## Supplementary Figures for "citrOgen: a synthesis-free polysaccharide and protein antigen-presentation to antibody-induction platform"

Wong and Sanchez-Garrdio et al.

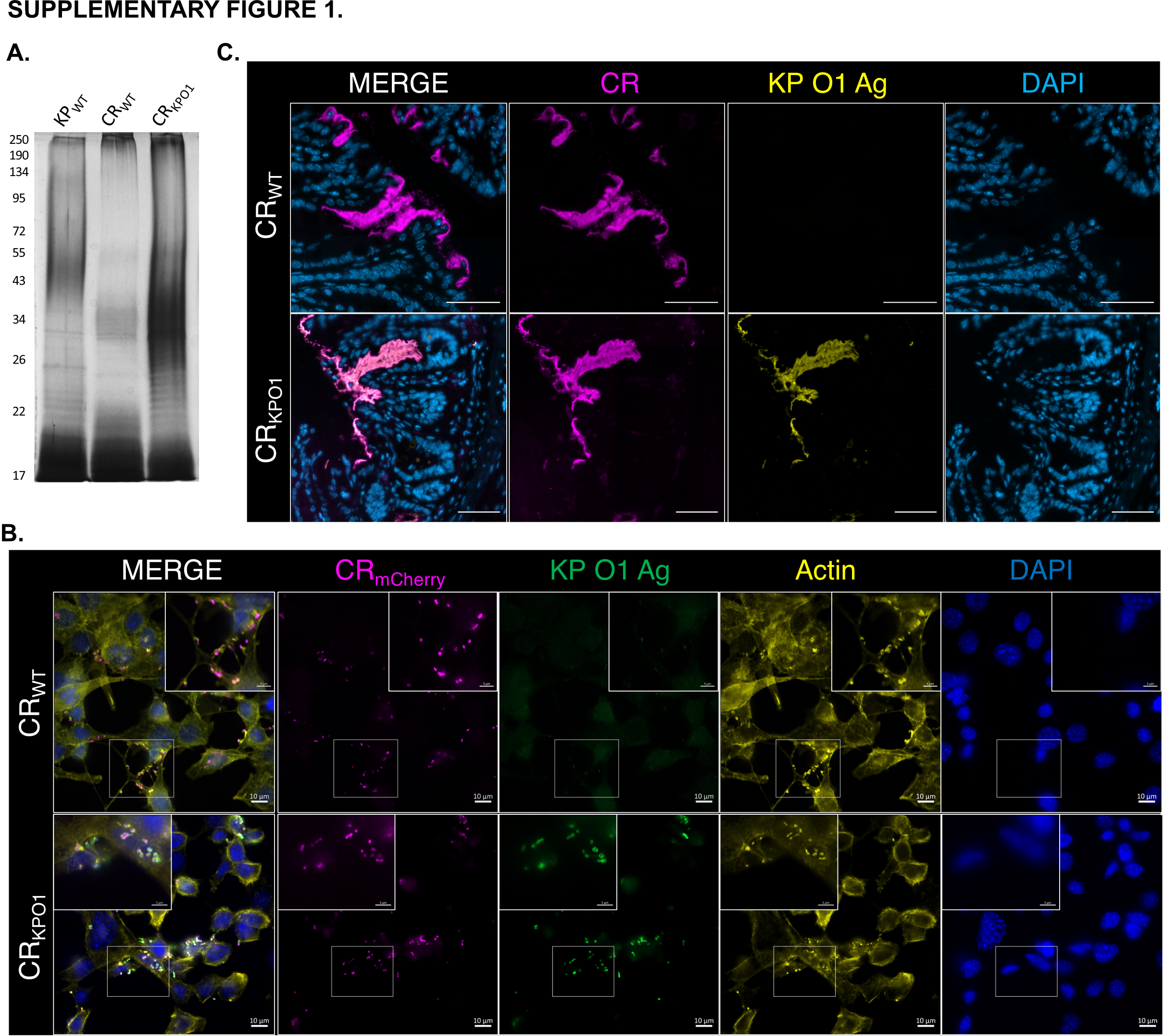

**Supplementary Figure 1 (related to Fig. 1). CR expressing a heterologous KP O-antigen (KPO1) triggers actin polymerisation *in vitro* but exhibits an *in vivo* colonisation defect.**

A. Silver stain SDS-PAGE gel of crude LPS samples from *K pneumoniae* wild-type (KP_WT_), *C. rodentium* wild-type (CR_WT_) and CR expressing KPO1 (CR_KPO1_).

B. Representative images of immunostaining of Swiss-3T3 cells infected with the indicated CR strains expressing pUltra-mCherry. CR_KPO1_ forms actin-rich pedestals (phalloidin F-actin stain) and is immunoreactive to an anti-KP O1 Ag monoclonal antibody (C13 mAb); nuclei were stained with 4',6-diamidino-2-phenylindole (DAPI). Scale bar = 10 µm.

C. Zoomed images of colonic sections at 8 dpi and stained for CR and KP O1 Ag; DAPI staining used as a structural marker. CR_WT_ has no detectable KP O1 Ag expression while CR_KPO1_ presents clear KP O1 Ag expression. Scale bar = 50 µm.

Images are representative of 2 (A and B) and 3 (C) biologically independent repeats.

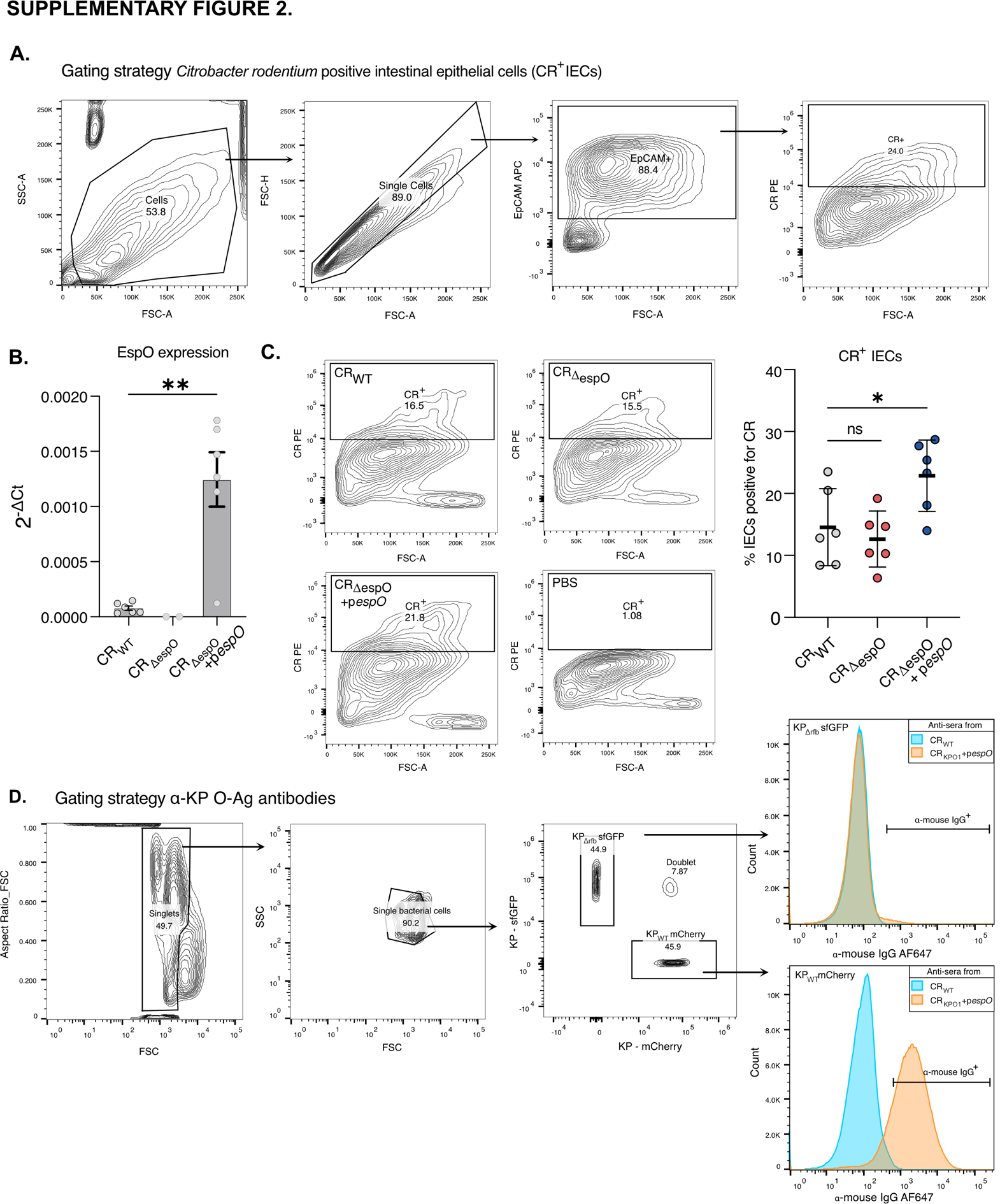

**Supplementary Figure 2 (related to Fig. 2). Overexpression of EspO in CR_KPO1_ increases attachment to IECs and induces specific anti-KP O1 Ag IgG responses.**

A. Gating strategy for the measurement of CR-positive (CR^+^) IECs (defined as EpCAM-positive).

B. *espO* expression was significantly higher in the presence p*espO* as determined by qRT-PCR in bacterial cultures grown statically in DMEM (T3SS-inducing conditions); CR_ΔespO_ was used as a negative control. Graph shows mean±SEM from 6 biological repeats; statistical significance was determined by paired t-test between CR_WT_ and CR_ΔespO_pespO_;_ ** p<0.01.

C. The proportion of CR^+^ IECs isolated from the distal colon is higher in mice infected with CR over-expressing *espO* from pACYC-*espO* (CR_ΔespO_+p*espO*); representative flow cytometry plots are shown on the left. Graph shows mean±SEM from 6 biological repeats; statistical significance was determined by repeated measured one-way ANOVA corrected with Dunnett’s multiple comparisons test; * p<0.05.

D. Gating strategy used to test the binding specificity of citrOgen-derived antibodies against KP O1-Ag using KP_WT_ expressing mCherry and KP_Δrfb_ expressing sfGFP. Representative histograms of the two populations are shown on the right, where the anti-sera from CR_KPO1_-infected mice is shown to bind KP_WT_ but not KP_Δrfb_, antisera from CR_WT_-infected mice were used as a negative control.

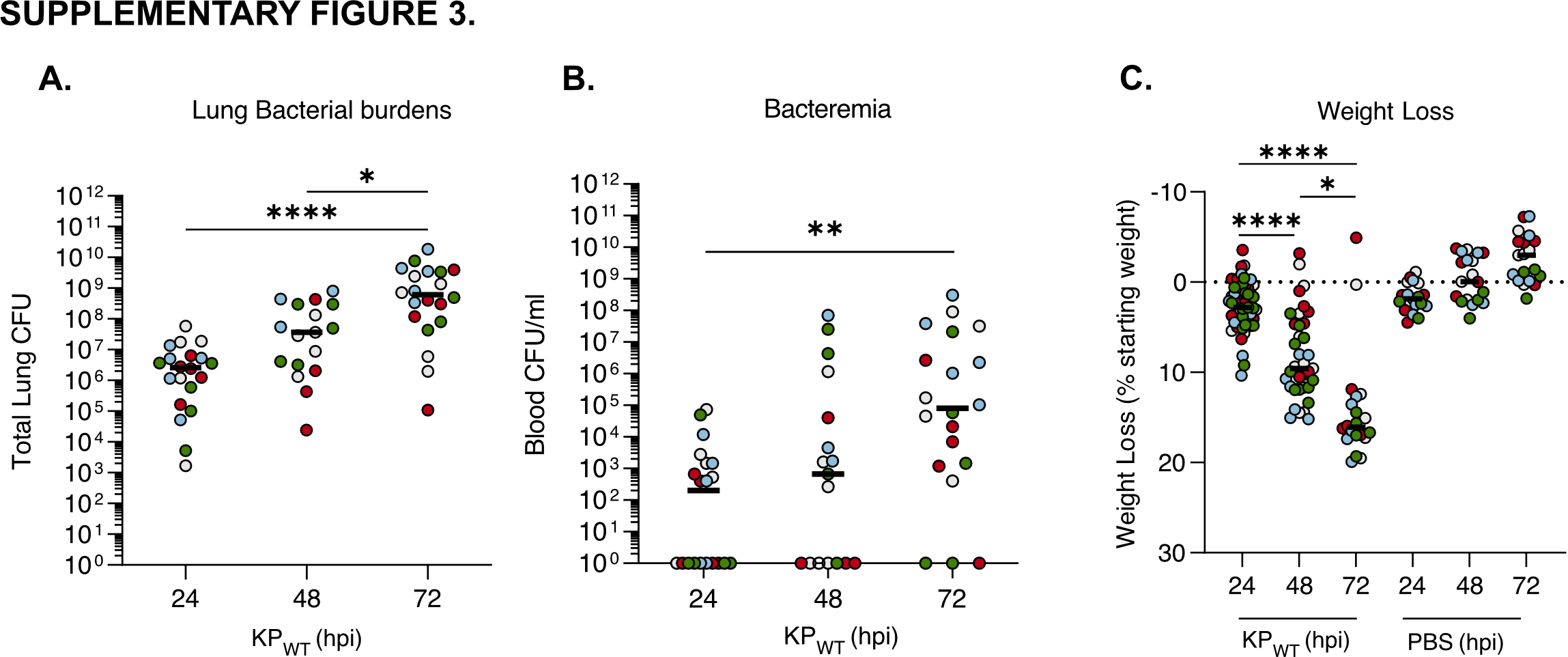

**Supplementary Figure 3. KP_WT_ lung infection (500CFU) of CD-1 mice results in consistent bacterial replication and disease progression.**

A. KP_WT_ burden in the lungs undergoes a stepwise daily increase, reaching significance by 72 hpi compared to 24 and 48 hpi.

B. The proportion of bacteraemic mice increases as the infection progresses; by 72 hpi 85% of the infected mice are bacteraemic.

C. KP_WT_ infection induces significant temporal weight loss; no weight loss was seen in the PBS mock-infected mice.

A-C. Graphs are representative of 4 biological repeats where each repeat is represented by a different colour. Statistical significance was determined by non-parametric Kruskal-Wallis test corrected with Dunn’s multiple comparisons test; in (C) significance was tested between the timepoints within a treatment group. * p<0.05; ** p<0.01; **** p<0.0001.

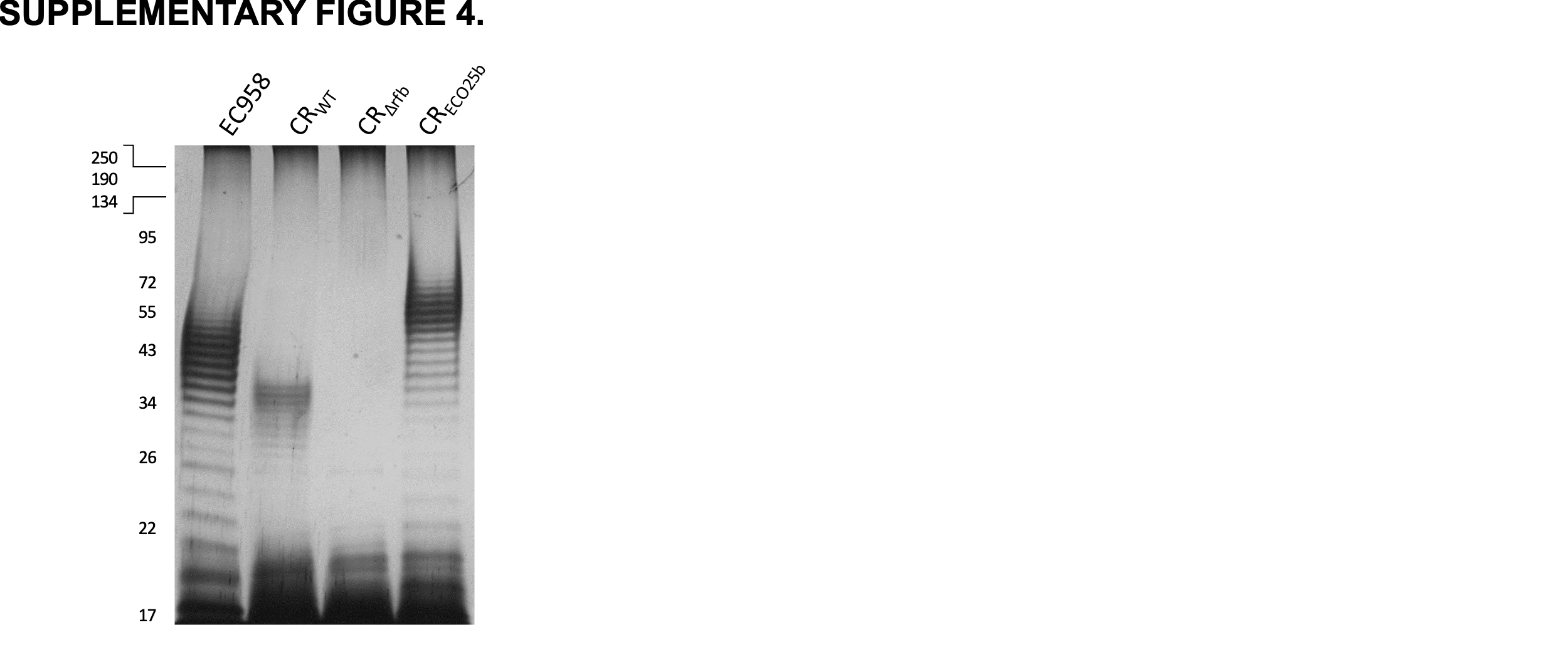

**Supplementary Figure 4 (related to Fig. 4). The citrOgen platform can be used to express O-antigens exported through the more common *wzy* pathway**

Silver stain SDS-PAGE gel of crude LPS samples from *E.coli* ST131 expressing O25b (EC958), *C. rodentium* wild-type (CR_WT_), CR lacking O-antigen expression (CR_Δrfb_) and CR expressing ST131 O25b (CR_ECO25b_).

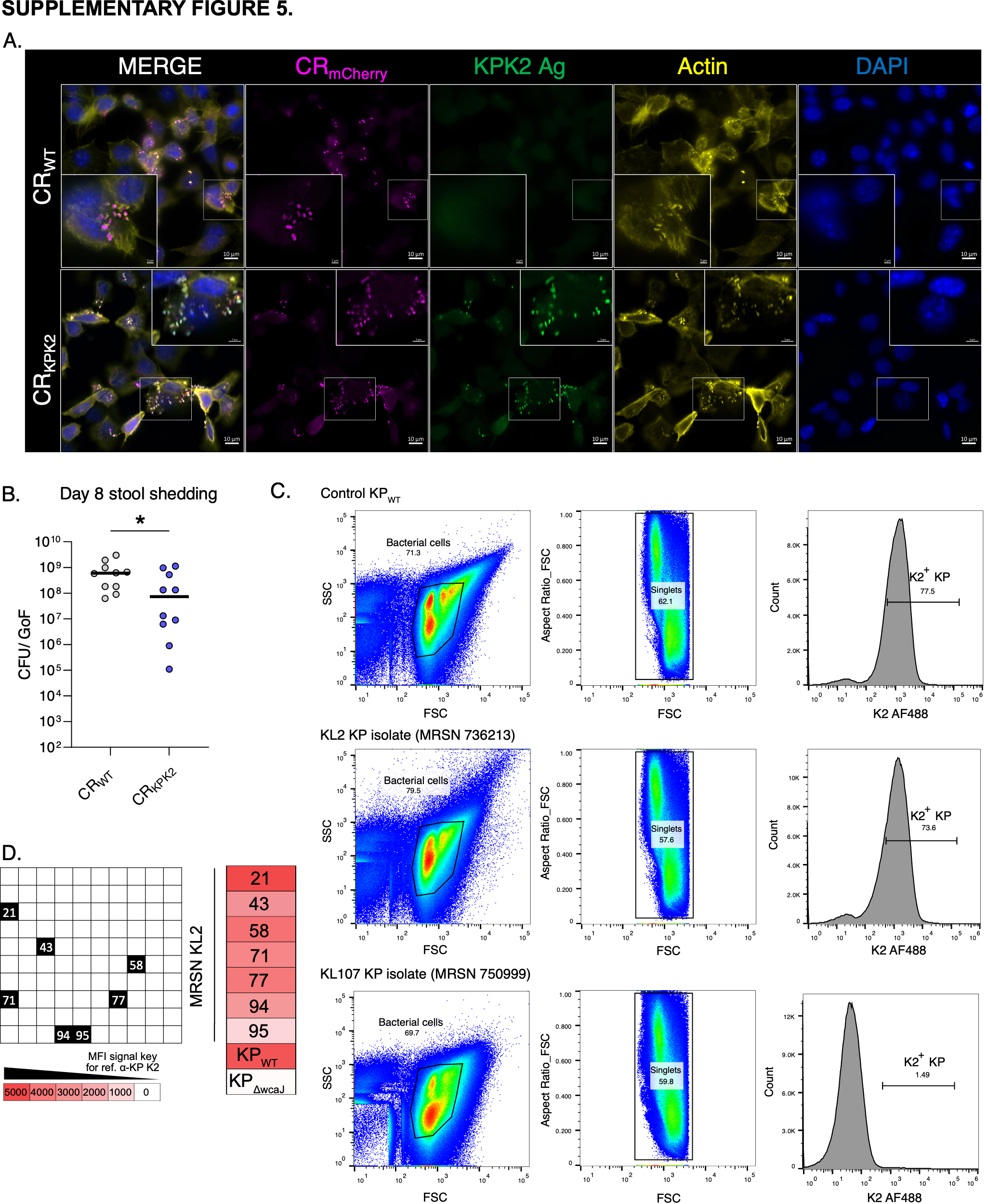

**Supplementary Figure 5 (related to Fig. 5). Anti-sera from CR_KPK2_-infected mice can be used for KP capsule serotyping.**

A. Representative images of Swiss-3T3 cells infected with the indicated CR strains expressing pUltra-mCherry and stained with anti-KP K2 antibody (SSI), phalloidin (F-actin stain) and DAPI (nucleic acid stain). CR_KPK2_ forms actin-rich pedestals while being immunoreactive to anti K2 KP antibodies. Scale bar = 10 µm. Images are representative of 2 biologically independent replicates.

B. At 8 dpi following oral gavage, faecal CFUs are significantly lower in CR_KPK2_ compared with CR_WT_ . Graph shows median of n=10 mice per group, 2 biological repeats. Statistical significance was determined using a lognormal Welch’s unpaired t test. *, P<0.05. Faecal CFUs for CR_WT_ are as in Supplementary Figure 6C. GoF, gram of faeces.

C. Representative flow cytometry dot plots obtained after staining the MRSN collection of 100 KP isolates with citrOgen-derived anti-KP K2 antibodies; KP_WT_ is used as a positive control. MRSN 736613 is shown to be a KL2-expressing isolate while MRSN 750999 is negative for KL2-staining, corroborating with the genomic analysis of these isolates predicted to be KL2 and KL107 respectively.

D. The reference anti-KP K2 antibody (SSI) was used to validate the results obtained for the KL2 strains from the collection, highlighted as numbered black squares in the schematic to the left, based on Fig 5E. KP_WT_ and the capsule-deficient KP_ΔwcaJ_ were used as controls. The boxes are coloured according to the strength of binding by the anti-KP K2 antibody, which was determined using the median fluorescence intensity obtained by flow cytometry analysis.

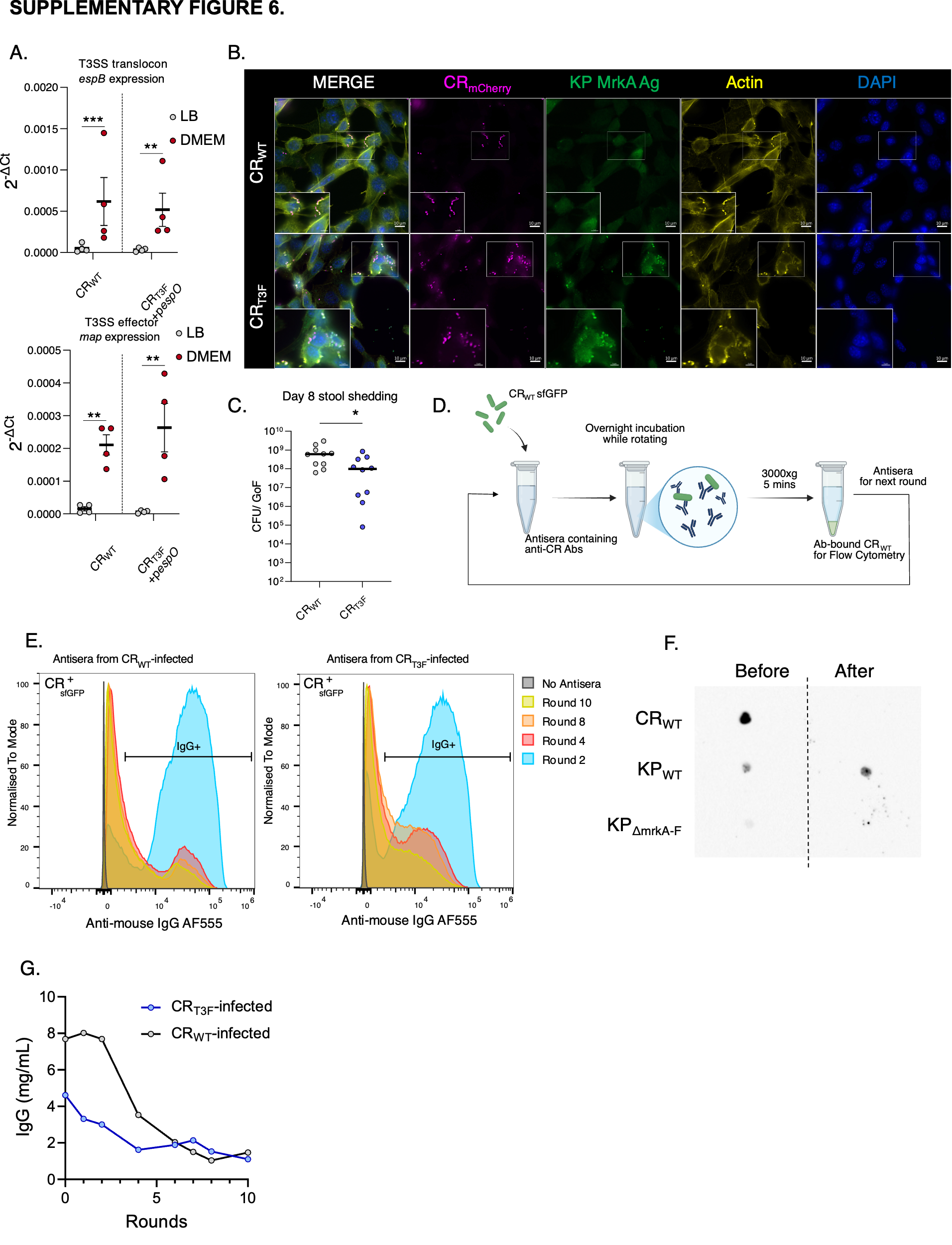

**Supplementary Figure 6 (related to Fig. 6). A complex protein Ag, KP T3F, expressed by citrOgen under the *map* promoter, is used to generate functional anti-KP T3F Abs.**

A. qRT-PCR was used to quantify the expression levels of the structural T3SS gene *espB* (top) and the T3SS effector *map* (bottom) in bacterial cultures of CR_WT_ and CR_T3F_ grown statically under T3SS suppressing (LB) and inducing (DMEM) conditions. T3SS-inducing conditions similarly induce *espB* and *map* transcription in CR_WT_ and CR_T3F_. Results are from 4 biological replicates; graphs show central tendency mean± SEM. Statistical significance was determined by ratio-paired t-tests comparing expression in LB and DMEM; ** p<0.01; *** p<0.001.

B. Representative images if Swiss 3T3 mouse fibroblasts infected with the indicated mCherry-tagged CR strains and immunostained with an Ab against MrkA, phalloidin (F-actin) and DAPI (nuclei). CR_T3F_ forms actin-rich pedestals to the same extent as CR_WT_ while expressing type 3 fimbriae (T3F), which also increased attachment to the glass coverslips. Scale bar=10 µm. Images are representative of 2 biological replicates.

C. At 8 dpi following oral gavage, faecal CFUs are significantly lower in CR_T3F_ compared with CR_WT_ . Graph shows median of n=10 mice per group from 2 biological repeats. Statistical significance was determined using a lognormal Welch’s unpaired t test. *, P<0.05. Faecal CFUs for CR_WT_ are as in Supplementary Figure 5B. GoF, gram of faeces.

D. Schematic representing the protocol used to pre-adsorb citrOgen-derived antisera using PFA-fixed CR expressing sfGFP (CR_WT_sfGFP) to remove CR_WT_ Abs.

E. Flow cytometry histograms of the antisera obtained from CR_WT_- (left) and CR_T3F_-infected (right) mice. which has undergone the indicated number of rounds of pre-adsorption against CR_WT_. While by round 2 of pre-adsorption most of the antisera still bound CR_WT_, by round 10 there is negligible binding to CR_WT_.

F. Dot-blot performed against the indicated bacterial strains with antisera from CR_T3F_-infected mice before and after 10 rounds of pre-adsorption normalised to the total IgG content; following pre-adsorption the antisera only bind KP_WT_.

G. Levels of total IgG in the antisera from CR_WT_ and CR_T3F_-infected mice assessed by ELISA, measured before, during and after the pre-adsorption process.
